# Recollection and prior knowledge recruit the left angular gyrus during recognition

**DOI:** 10.1101/561910

**Authors:** B. Bellana, N. Ladyka-Wojcik, S. Lahan, M. Moscovitch, C. L. Grady

**Author notes:** **Authors’ Note**: Correspondence can be addressed to Buddhika Bellana, Cheryl Grady, or Morris Moscovitch. Author contributions*: B.B., C.L.G. & M.M. generated hypotheses and designed the experiment; B.B., N.L.W, and S.L. collected the data; B.B. analyzed the data. All authors contributed to writing and approved of the final version of the submitted manuscript.

## Abstract

The human angular gyrus (AG) is implicated in recollection, or our ability to retrieve detailed memory content from a specific study episode. Parallel work also highlights a key role of the AG in the representation of general knowledge and semantics. How these two lines of research converge remains unclear. The present fMRI experiment used a remember-know paradigm with famous and non-famous faces to test whether activity in the AG could be modulated by both task-specific recollection and general prior knowledge in the same participants. Increased BOLD activity in the left AG was observed during both recollection in the absence of prior knowledge (i.e., recollected > non-recollected or correctly rejected non-famous faces) and when prior knowledge was accessed in the absence of recollection (i.e., famous > non-famous correct rejections). This pattern was unique to the left AG, and was not present in any other regions of the lateral inferior parietal lobe. Furthermore, the response profile of the left AG was consistent with accounts of recollection strength. Recollection-related activity was greater for faces with longer exposures at encoding than those with shorter exposures and was greater for stimuli with prior knowledge than those without, despite prior knowledge being incidental to the recognition decision. Therefore, the left AG is recruited during the access of both task-specific recollection and general prior knowledge, with greater activity as the amount of retrieved information increases, irrespective of its episodic or semantic nature.

**Significance Statement:** The human angular gyrus (AG) is often implicated in our ability to remember past events. A separate line of research examining our ability to represent general knowledge has also highlighted the AG as a core region of interest. To reconcile these separate views of AG function, we used fMRI to test whether the human left AG was sensitive to remembering details from a specific study episode (i.e., recollection) or more general prior knowledge, within the same participants. Overall, activity in the left AG was sensitive to *both* recollection and prior knowledge, suggesting any complete functional account of the left AG during retrieval must consider its sensitivity to both kinds of mnemonic representations.

## INTRODUCTION

The angular gyrus (AG) is reliably implicated in our ability to remember past events (Cabeza, Ciaramelli, & Moscovitch, 2012; Levy, 2012; Rugg & King, 2017; Sestieri, Shulman, & Corbetta, 2017; Shimamura, 2014; Wagner, Shannon, Kahn, & Buckner, 2005). Evidence from neuroimaging suggests that the AG is a critical node in a distributed network of brain regions that are reliably engaged during recollection (Frithsen & Miller, 2014; Rugg & Vilberg, 2013). Activity in the AG tracks how precisely a representation in memory matches the original properties of the encoded stimulus (Richter, Cooper, Bays, & Simons, 2016), such that physical stimulus properties can be decoded from memory-related activity (Kuhl & Chun, 2014; Lee & Kuhl, 2016; St-Laurent, Abdi, & Buchsbaum, 2015). Access to these precise representations can be temporarily modulated using non-invasive stimulation targeting the AG, further supporting its direct involvement in recollection (Bonnici, Cheke, Green, FitzGerald & Simons, 2018; Nilakantan, Bridge, Gagnon, Vanhaerents, & Voss, 2017; Thakral, Madore, & Schacter, 2017; Wang et al., 2014; Yazar, Bergström, & Simons, 2017).

The left AG, however, is not exclusively involved in episodic memory and recollection. Accessing semantic memory, or the general knowledge abstracted over multiple past experiences, reliably recruits the left AG (Binder & Desai, 2011; Binder, Desai, Graves, & Conant, 2009; Kim, 2016; Price, 2010; Seghier, 2013), and access to semantics can be modulated with non-invasive brain stimulation targeting this region (Capotosto et al., 2016; Price, Peelle, Bonner, Grossman, & Hamilton, 2016).

Overall, evidence suggests that the left AG is implicated in the simultaneous representation of sensory-rich, idiosyncratic details from specific past experiences (i.e., recollection) and general knowledge derived across multiple past episodes. These two types of information are often considered distinct (McClelland, McNaughton, & O’Reilly, 1995; Squire, 1986; Tulving, 1972; Winocur, Moscovitch, & Bontempi, 2010), which may contribute to the paucity of attention that has been given to their co-occurrence in current theoretical models of AG function (for recent discussion, see Ramanan et al., 2017; Rugg & King, 2017; Seghier, 2013). Their co-occurrence in the AG in particular is consistent with the idea that this region acts as an integrator (Bonner, Peelle, Cook, & Grossman, 2013; Bonnici, Richter, Yazar, & Simons, 2016; Price, Bonner, & Grossman, 2015; Ramanan et al., 2017), potentially able to extend beyond the sensory domain to combine mnemonic content across specific recollections and general prior knowledge.

Despite evidence suggesting the common recruitment of the left AG when accessing specific recollections and general prior knowledge, a direct test of this hypothesis, within the same individuals and experimental paradigm, has not yet been conducted. In the present fMRI experiment, participants’ recognition memory for faces of famous and non-famous individuals was assessed using a Remember-Know paradigm. Famous faces are associated with multiple past episodes, and semantic person knowledge is rapidly accessed upon presentation of the face (Bruce & Young, 1986; Ramon & Gobbini, 2017), thus providing an ideal stimulus class for eliciting prior knowledge. Importantly, the presence or absence of prior knowledge is orthogonal to any recognition decision in the context of a standard Remember-Know paradigm (Tulving, 1985), allowing us to independently capture effects of prior knowledge and experiment-specific recollection. Additionally, we manipulated encoding duration (Leiker & Johnson, 2014; Vilberg & Rugg, 2009a, 2009b) to determine whether recollection-related activity in the left AG was sensitive to differences in the amount of recollected content (Hutchinson et al., 2014; Rugg & King, 2017). Using this paradigm, we were able to directly test the independent effects of recollection and prior knowledge on activity in the left AG and the surrounding inferior parietal lobe. We also explored whether activity in the left AG tracked recollection and prior knowledge simultaneously, consistent with its role in integration. Specifically, we predicted that the left AG should track both recollection and prior knowledge, thereby bridging the episodic and semantic memory literatures and providing evidence for a general role in tracking the amount of retrieved mnemonic content.

## MATERIALS & METHODS

### Participants

Twenty-nine young adults between 19-30 years of age participated in the experiment. Participants were recruited from the University of Toronto and surrounding areas, and were compensated a total of $50 with an average testing session lasting 2.5 hours. Five participants did not meet our inclusion criteria for the final sample and were excluded from subsequent analyses (1 did not recognize at least 50% of famous stimuli, and 4 had excessive movement or drowsiness during scanning). Our final sample included 24 participants (years of age: *M* = 22.5, *SD* = 2.4; years of education: *M* = 16.5, *SD* = 1.9; n_female_ = 11). All participants were screened to be healthy, right-handed and free of health problems/medications, psychiatric or neural diagnoses or vascular disease, that may influence cognitive functioning or brain activity. The study was approved by the Research Ethics Board at the University of Toronto and informed consent was given by each participant before participating.

### Stimuli

Images were obtained from the Internet using Google image search. We included 128 highly recognizable faces and 128 age/race matched non-famous counterparts as stimuli, for a total of 256 faces (for further details, see Bellana, Mansour, Ladyka-Wojcik, Grady & Moscovitch, 2019). Both famous and non-famous stimulus pools were balanced in terms of sex (n_male_ = 64, n_female_ = 64, for both famous and non-famous pools) and were all neutral to slightly positive in expression. All faces were manually centered and cropped from the full image using an oval frame in Adobe Photoshop and resized to 475×595 pixels. Images were set to black and white and their luminance was matched using SHINE toolbox (Willenbockel et al., 2010). A scrambled version of each face was also generated using custom scripts in Matlab (MathWorks, Natick, MA, USA), where each image was divided into 5-pixel clusters and then randomly shuffled. These scrambled images were used for null trials in the fMRI experiment.

### Experimental Procedure

Overall, participants underwent an eyes-open resting state scan (7 minutes, 8 seconds), then performed 4 experimental recognition memory blocks (7 minutes, 8 seconds each), followed by a structural scan (7 minutes, 10 seconds), 4 experimental recognition memory blocks, and a final eyes-open resting state scan for a total time of approximately 80 minutes in the scanner. After scanning, participants underwent a final surprise self-paced delayed recognition test outside of the scanner (*Mean duration of post-scan test* = 20 minutes, 45 seconds). The total session, including consent, instructions, practice, scanning preparation and debriefing, was completed within 2.5 hours.

In the scanner, we assessed memory using a Remember-Know paradigm on a total of 128 famous and 128 non-famous faces (96 targets, 32 lures each). For a schematic overview of the experiment, see Figure 1A. Remember-Know is a commonly used recognition memory paradigm to measure recollection and familiarity (Tulving, 1985; Yonelinas, 2002). In addition to making an old-new decision during recognition, participants introspect about the subjective quality of their recognition. Participants select the Remember option (R) if their recognition is accompanied by recall of contextual information from the study episode. Know (K) is selected if recognition is not accompanied by any contextual information from study. Behavioural estimates of recollection and familiarity were calculated using the method from Yonelinas & Jacoby (1995), while adjusting for false alarms. Recollection was defined as R_hit_ − R_fa_, or the proportion of studied trials that received an R response (i.e., *hit*) minus the proportion of new trials that received an R response (i.e., false alarms; *fa*). Familiarity was defined as K_hit_/(1 – R_hit_) − K_fa_/(1 – R_fa_), or the number of old trials that received a Know response divided by the number of old trials that did not receive a Remember response and correcting for false alarms by subtracting the same ratio calculated using new trials.

**Figure 1.**
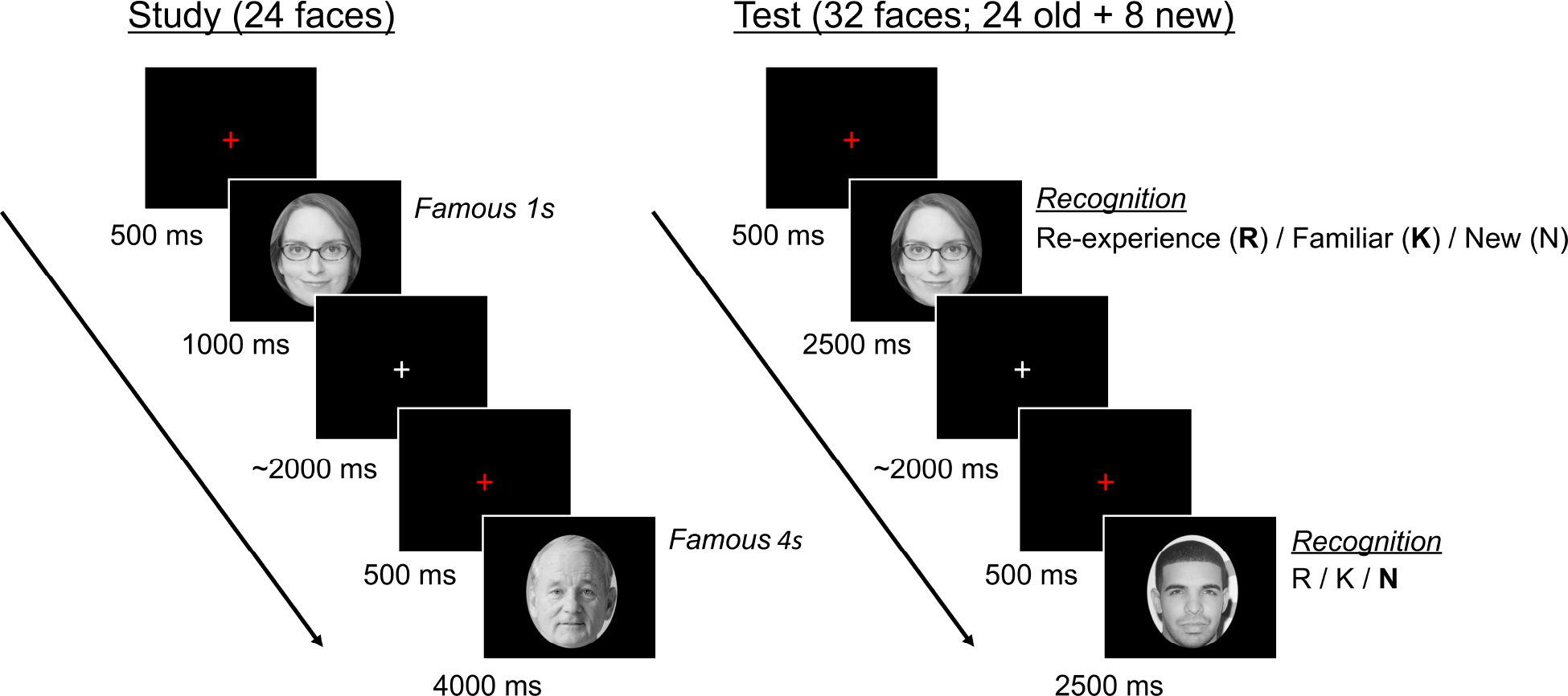
Schematic example of one in-scan study-test block. For details, see Procedure.

The experimental paradigm consisted of 8 blocks, each of which contained a study and recognition phase (Figure 1). Prior knowledge was manipulated at the block level such that 4 of the blocks were entirely composed of famous stimuli and the remaining 4 blocks of non-famous stimuli, in a fixed interleaved order which was counterbalanced across participants. During each encoding phase, participants explicitly studied a pseudorandom sequence of 24 face trials. Each trial began with a red fixation cross presented for 500ms, followed by a face stimulus presented centrally onscreen. Half of the face stimuli were presented for 1s and half for 4s to provide a manipulation of encoding duration, followed by an inter-trial interval ranging between 1500 to 2750ms (*M* = ~2s). Participants were asked to study each face in as much detail as possible as their memory would be tested immediately after each study phase. No explicit response was required during study, though eye-movement was monitored by the experimenter via eye-tracking to ensure participants were fixating on the faces. Participants were also instructed that each block was independent from the others and contained unique face stimuli. In addition to the target face trials, 12 additional null trials were also interspersed during the study phase. Null trials consisted of scrambled face stimuli from the experiment, 6 of which were presented for 1s and 6 for 4s. These trials were not to be encoded, but were collected to serve as a potential baseline condition.

The first recognition trial was presented 20s after the termination of the last study trial. During each recognition phase, participants were presented with a pseudorandom sequence of 32 faces, 24 of which were from the previous study list and 8 were novel lures (i.e., old/new ratio of 75/25). Each trial began with a red fixation cross presented for 500ms, followed by the face stimulus presented centrally onscreen. Each face was presented for 2.5s during which time participants were required to make a Remember (R), Know (K), or New (N) response using a fMRI compatible response box. Participants used their right hand to make responses, with the index and middle finger being used to respond R or K and the ring finger, to respond N. The mapping of index and middle finger to the R and K response option was counterbalanced across participants. In addition to the studied target and novel lure recognition trials, 12 additional null scramble trials were also interspersed during the recognition phase. Participants were asked to press any of the three response keys for these trials, with no demands on recognition.

After the 8 study-test blocks were completed, participants exited the scanner and were briefly interviewed regarding their experience and memory strategies. Lastly, participants were asked to perform a surprise delayed recognition memory test. All 256 faces from the previous 8 blocks of the experiment (i.e., both studied targets and novel lures) in addition to 32 entirely novel unstudied faces (16 famous and 16 non-famous) were presented in a random order during the delayed test. Trials were self-paced and participants responded with the index fingers of both hands using the buttons Q or P on the keyboard with response mapping counterbalanced across participants. Each trial began with a 500ms red fixation cross, followed by the presentation of a face in the centre of the screen. Participants were first required to indicate whether the face was old (i.e., a target or lure previously seen in the experiment) or new (i.e., entirely novel, not seen in the experiment). Next, participants were asked to indicate whether they believed the face was of a famous or non-famous individual, based on their personal experience. For faces judged as famous, participants were then required to indicate how much they knew about the individual on a scale of 1 (very little) to 5 (a great deal) using the number pad on the keyboard. Participants then indicated whether or not they could name the famous individual via a yes or no button response.

### fMRI data acquisition

MRI images were acquired using a Siemens Prisma 3T scanner with a 32-channel head coil at the Toronto Neuroimaging (ToNI) centre located at the University of Toronto. Structural MRI images were collected using a T1-weighted high-resolution scan with a standard 3-dimensional magnetization-prepared rapid-acquisition gradient echo (MPRAGE) pulse sequence [160 slices; field of view (FOV) = 256 × 256 mm; 1 mm isotropic resolution; echo time (TE) = 2.4 ms; repetition time (TR) = 2000 ms; flip angle = 9°; for a total duration of 430 s]. For the functional MRI images (both task and rest), blood oxygenation level-dependent (BOLD) signal was measured using a T2-weighted multiband echo planar imaging (EPI) acquisition procedure [39 slices; FOV = 216 × 216 mm; 3 mm isotropic resolution; TE = 31 ms; TR = 1000 ms; flip angle = 45°; multiband factor = 3, with a total of 428 volumes collected]. Head movements were limited by inserting soft cushions into the head coil. Eye movements were monitored using an in-scan MRI compatible EyeLink 1000 plus (SR research, Ltd.). Visual stimuli were presented by E-Prime software (version 2, Psychology Software Tools, Inc.), presented on an MRI-compatible LCD screen and viewed with a mirror mounted on the head coil. Responses were collected with an MRI-compatible response box.

### Data Preprocessing

Preprocessing of MRI images was conducted using a combination of functions from AFNI (Cox, 1996; https://afni.nimh.nih.gov/) and FSL (Smith et al., 2004; https://fsl.fmrib.ox.ac.uk/fsl/fslwiki). The pipeline included the following steps: DICOM to NII (AFNI: Dimon), spatial realignment (AFNI: 3dvolreg), co-registration (FSL: epi_reg), subject-specific tissue segmentation (FSL: FAST) and spatial normalization to MNI space (FSL: FLIRT). The final functional images were resampled to 2 mm isotropic voxels (AFNI: 3dresample), spatially smoothed using a Gaussian kernel with the full-width at half maximum of 5 mm (AFNI: 3dmerge). An additional motion scrubbing procedure was added to the end of our preprocessing pipeline (Campbell et al., 2013). Using a conservative multivariate technique, volumes that were outliers in both the six rigid-body motion parameter estimates and BOLD signal intensity were removed, and replaced by interpolating the BOLD signal across neighbouring volumes. Motion scrubbing further minimizes any effects of motion-induced spikes on the BOLD signal, over and beyond standard motion regression (which was included in the subsequent analysis step), without leaving sharp discontinuities due to the removal of outlier volumes (for details, see Campbell et al., 2013). Images were manually inspected throughout the preprocessing pipeline to assure data quality.

### Defining regions of interest

Since our hypotheses were specific to the left AG, a region of interest (ROI)-based approach at the group-level was most appropriate. Considering the structural and functional heterogeneity of the inferior parietal lobe (Caspers et al., 2006, 2013; Nelson et al., 2010), we chose to use a fine-grained ROI definition based on cytoarchitectural probability maps from the Jülich Histological Atlas (Eickhoff et al., 2005) as implemented in FSL. The inferior parietal lobe consists of the supramarginal gyrus (anteriorly) and the angular gyrus (posteriorly), which roughly correspond to Brodmann’s area 40 and 39. More recent studies of cytoarchitecture suggest that the inferior parietal lobe consists of 7 separable regions, listed in the anterior to posterior direction: PFop, PFt, PFcm, PF, PFm, PGa and PGp (see Caspers et al., 2006, 2011, 2013). Tissue probability maps for these 7 regions across both left and right hemispheres were selected from the Jülich Histological Atlas (Eickhoff et al., 2005) and thresholded at 40% as a conservative approach to form our ROIs (see Figure 4A, and Extended Data: Figure 4-1). These maps were further exclusively masked, from posterior to anterior, to ensure all ROIs were non-overlapping. From these masks, regions PGa and PGp are the most posterior and fall within the vicinity of the AG. Notably, the more posterior PGp has robust structural connectivity with the hippocampus while the PGa does not (Uddin et al., 2010). Furthermore, a Neurosynth (Yarkoni et al., 2011) reverse inference meta-analytic map based on the term “episodic memory” (http://neurosynth.org/analyses/terms/episodic%20memory/; derived from 270 studies, FDR corrected at p < 0.01, 2 mm isotropic voxel resolution) produced a total of 350 voxels overlapping with the left AG (i.e., defined as a combined mask of the left PGa and PGp), such that 68% of these fell within the PGp and the remaining 32% within the PGa. For additional context, only 37 voxels associated with episodic memory fell in subregions anterior to the left PGa. Therefore, we selected the left PGp as our primary focus for subsequent analyses.

### fMRI Analysis

A general linear model (GLM), via AFNI’s 3dDeconvolve, with a gamma response function was used to estimate BOLD activity for each trial type per participant during retrieval. The GLM included 12 regressors of interest: 1) recollected (i.e., remember response) famous trials with 4s exposure at encoding [*M* = 37.7 trials, *SD* = 6.1], 2) recollected famous trials with 1s exposure at encoding [*M* = 35.2 trials, *SD* = 6.3], 3) recollected non-famous trials with 4s exposure at encoding [*M* = 30.8 trials, *SD* = 6.1], 4) recollected non-famous trials with 1s exposure at encoding [*M* = 24.5 trials, *SD* = 7.9], 5) non-recollected (i.e., know response or miss) famous trials with 4s exposure at encoding [*M* = 8.2 trials, *SD* = 6.1], 6) non-recollected famous trials with 1s exposure at encoding [*M* = 10.7 trials, *SD* = 6.4], 7) non-recollected non-famous trials with 4s exposure at encoding [*M* = 15.5 trials, *SD* = 6.3], 8) non-recollected non-famous trials with 1s exposure at encoding [*M* = 15.5 trials, *SD* = 6.3], 9) correctly rejected new famous trials [*M* = 27.5 trials, *SD* = 4.0], 10) correctly rejected new non-famous trials [*M* = 25.4 trials, *SD* = 5.0], 11) scrambled null trials [96 trials], and 12) button presses across all trials [352 trials]. Note that regressors 1-4 include only accurately recollected trials by definition. Averaged timecourses from subject-specific white matter and cerebrospinal fluid masks (thresholded at 90% probability) alongside 6 rigid-body motion parameters were also modelled as regressors of no interest. Furthermore, 1-4^th^ order polynomial trends were included in the model to account for scanner drift and other extraneous changes in the signal over time. To ensure that we were manipulating prior knowledge accurately within our participants we only modeled trials that were accurately judged as famous or non-famous by our participants according to post-scan fame judgements (*mean number of omitted trials* = 10/256, or 4%). To account for the low trial counts in the non-recollected famous conditions, estimates were collapsed over duration in all applicable subsequent group analyses. Relevant voxelwise beta estimates were then averaged within each ROI, per participant, and then submitted to subsequent group level analyses of variance (ANOVA), pairwise t-tests and trend analyses. To further supplement our gamma response-based model and to take advantage of our multiband imaging sequence, we conducted an additional GLM using a finite impulse response (FIR) approach to visualize the timecourse of activity in the PGp. FIRs were estimated using an 18 parameter tent function in AFNI’s 3dDeconvolve (*TENTzero*) from 1 second before stimulus onset to 17 seconds after, and were thus aligned to our 1 second TR grid. The results from this model were used for visualization only. Whole-brain results were visualized using BrainNet Viewer (Xia, Wang, & He, 2013; http://www.nitrc.org/projects/bnv/).

## RESULTS

### Behavioural results

#### Prior knowledge and encoding duration improve overall recognition

Adjusted recognition accuracy was defined as the proportion of accurately recognized targets after subtracting the proportion of false alarms on novel lures (hits – false alarms), per participant. This measure characterizes overall recognition while collapsing over subjective distinctions in recognition quality, namely, Remember or Know. The effect of prior knowledge and encoding duration on adjusted recognition accuracy was measured using a 2 (prior knowledge: famous, non-famous) × 2 (encoding duration: 1s, 4s) repeated measures ANOVA. Group-averaged accuracy across conditions is presented in Figure 2A. Significant main effects of both prior knowledge [*F*(1,23) = 42.59, *p* < 0.0001, *partial η^2^* = .65] and encoding duration [*F*(1,23) = 48.23, *p* < 0.0001, *partial η^2^* = .68] were observed, indicating that the presence of prior knowledge and a longer encoding opportunity during study both improved subsequent recognition accuracy. A significant interaction between prior knowledge and encoding duration was also observed [*F*(1,23) = 8.00, *p* = 0.01, *partial η^2^* = .26]. Simple effects using pairwise t-tests demonstrated that the benefit of longer encoding duration was most prominent during encoding of non-famous (*M*_1s_ = .635, *M*_4s_= .714, *SD_diff_* = 0.060; *t*(23) = 6.50, *p* < .0001, *Cohen’s d* = 1.33) relative to famous faces (*M*_1s_ = .834, *M*_4s_= .874, *SD_diff_* = 0.048; *t*(23) = 4.11, *p* = .0004, *d* = .84).

**Figure 2.**
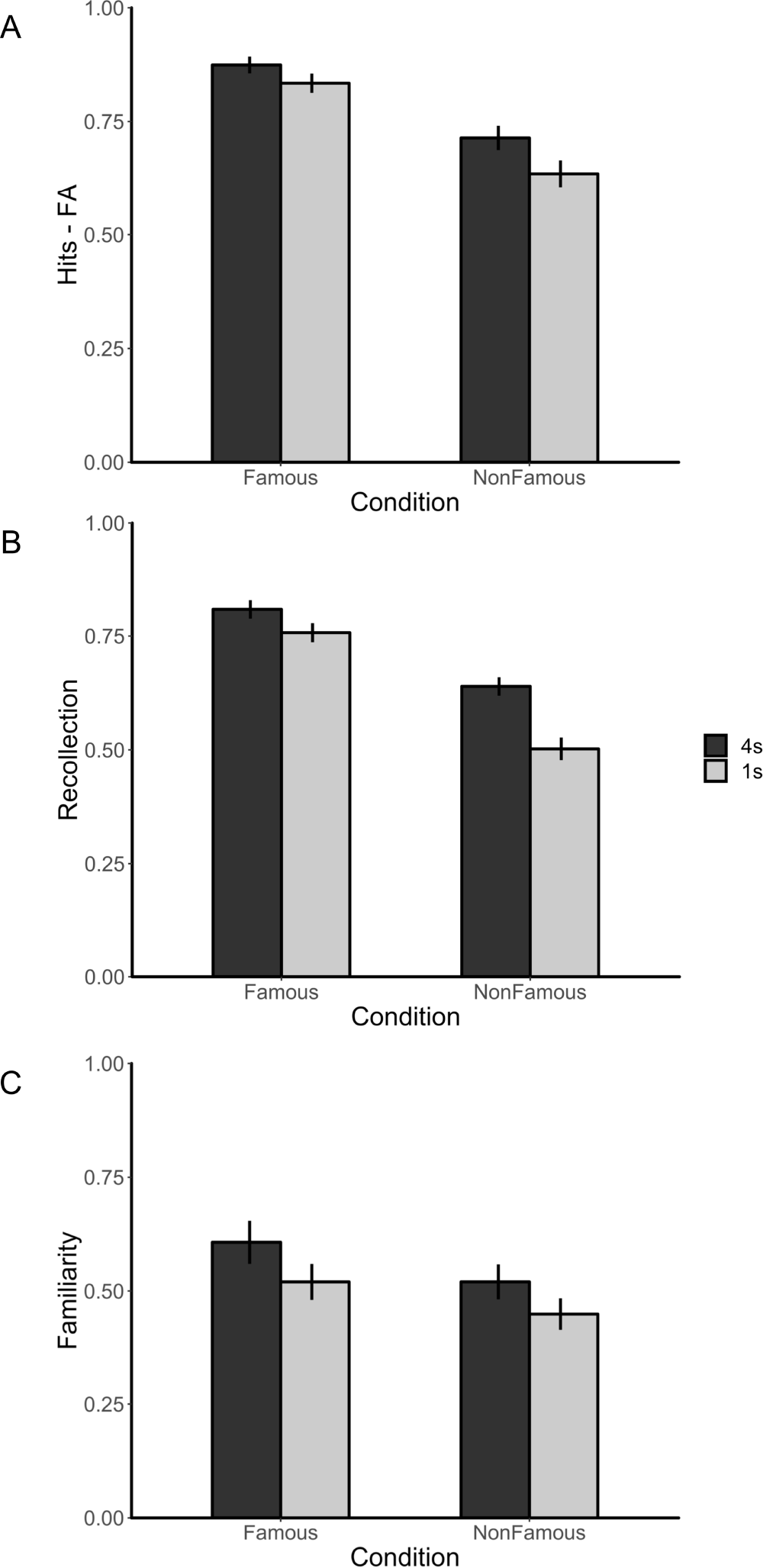
Behavioural performance. Effect of prior knowledge (famous, non-famous) and encoding duration (1s, 4s) on A) overall recognition (i.e., hits – false alarms), B) recollection (i.e., proportion of Remember hits – proportion Remember false alarms), and C) familiarity (i.e., proportion of all Non-Remembered target trials with Know responses – proportion of all Non-Remembered new trials with Know responses). Darker bars represent the 4s encoding duration, while lighter bars represent 1s. Error bars represent standard error of the mean. For additional behavioural results, see Extended Data: Figure 2-1.

#### Prior knowledge and encoding duration increase likelihood of recollection

The subjective quality of recognition was then examined by submitting the estimates of recollection (R) and familiarity (F) to a 2 (prior knowledge: famous, non-famous) × 2 (encoding duration: 1s, 4s) × 2 (recognition type: R, F) repeated measures ANOVA. Group-averaged estimates for recollection and familiarity across conditions are presented in Figure 2B-C. Main effects of prior knowledge [*F*(1,23) = 39.79, *p* = 0.0002, *partial η^2^* = .63], encoding duration [*F*(1,23) = 13.67, *p* = 0.001, *partial η^2^* = .37] and recognition type [*F*(1,23) = 7.42, *p* = 0.012, *partial η^2^* = .24] reached significance. Recognition type, however, interacted with prior knowledge [*F*(1,23) = 7.93, *p* = 0.0099, *partial η^2^* = .26], whereas other interactions failed to reach significance (all *p*s > .3). The interaction between prior knowledge and recognition type was decomposed using paired t-tests comparing the effect of prior knowledge for recollection and familiarity separately, while collapsing over encoding duration. Simple effects indicate that prior knowledge had a differential impact on estimates of recollection (*M*_Famous_ = .783, *M*_Non-famous_ = .571, *SD_diff_* = 0.222; *t*(23) = 12.32, *p* < .0001, *Cohen’s d* = .96), with no significant effect on familiarity (*M*_Famous_ = .564, *M*_Non-famous_ = .485, *SD_diff_* = 0.323; *t*(23) = 1.81, *p* = .083, *Cohen’s d* = .24). The benefit of prior knowledge on recollection estimates has been reported previously using this paradigm (see Bellana et al., 2019).

### fMRI results

#### Recollection and prior knowledge modulate activity in the left AG

The primary question of interest was to determine whether activity in the left AG could be modulated both by task-specific recollection and prior knowledge in the same participants with the same paradigm. We, therefore, submitted participant-specific parameter estimates averaged across all voxels in the left PGp to a 2 (prior knowledge: famous, non-famous) × 2 (recollection: recollected, correct rejection) repeated measures ANOVA. Parameter estimates for recollection trials were averaged across those presented for 4s and 1s at study to first test the effect of recollection regardless of encoding duration. Timecourses of activity based on FIR models of left PGp activity across all trials of interest are presented in Figure 3A-B. Group-averaged parameter estimates across conditions are presented in Figure 3C. Main effects of recollection [*F*(1,23) = 6.26, *p* = 0.02, *partial η^2^* = .21] and prior knowledge [*F*(1,23) = 13.03, *p* = 0.001, *partial η^2^* = .36] were observed. The interaction between recollection and prior knowledge failed to reach significance [*F*(1,23) < 1]. Overall, this suggests that the left AG 1) responds to recollection irrespective of whether the face was previously known or novel, and 2) responds to prior knowledge irrespective of whether the trial was accurately recollected or correctly rejected.

**Figure 3.**
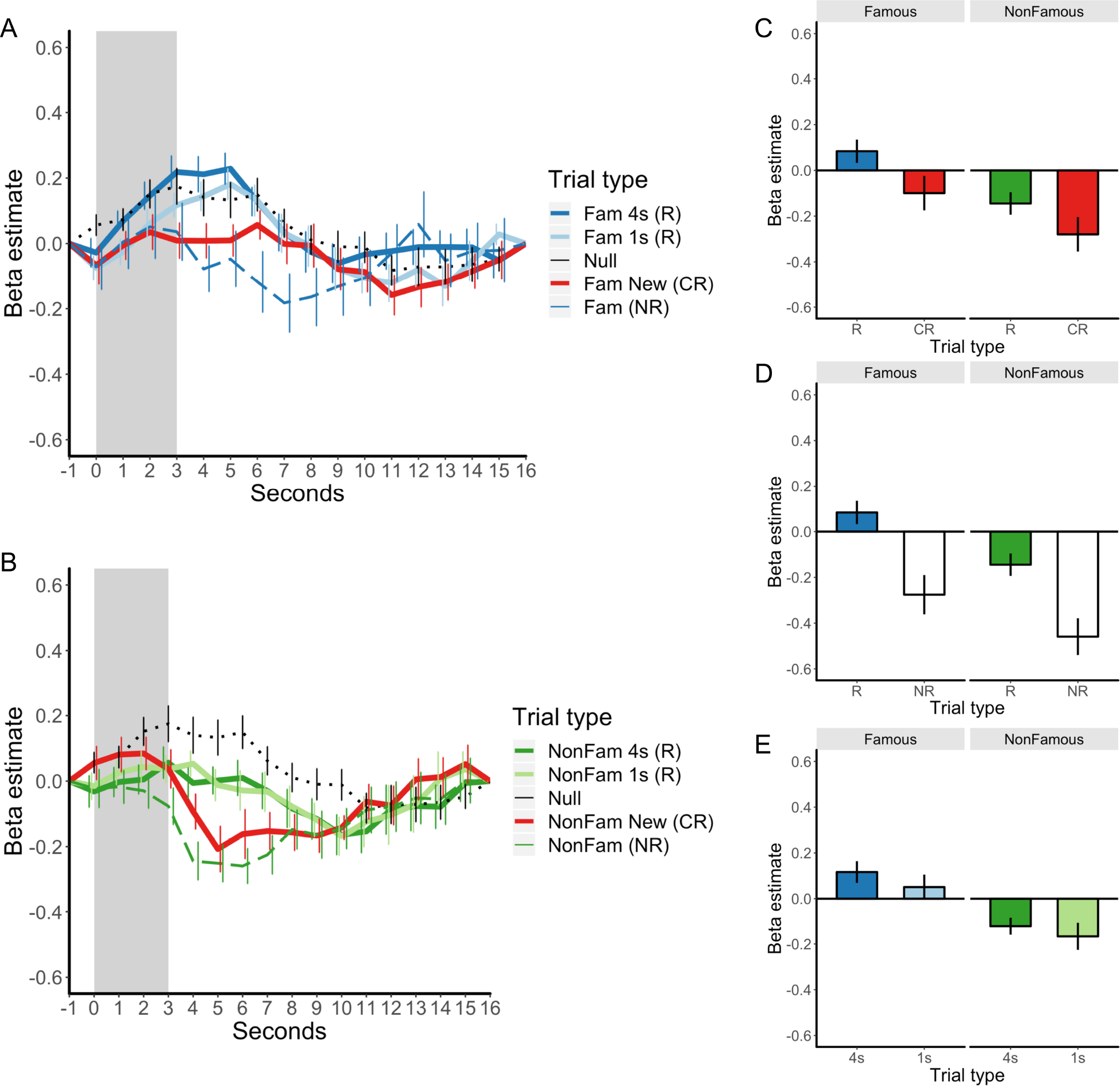
Time-course of left PGp activity based on a finite impulse response model, visualizing trial-average activity patterns for A) famous (blue) and B) non-famous (green) trials separately. New famous and non-famous trials are plotted in red. Dotted line in black represents the time-course for null trials. *Abbreviations*: R: Recollected target (i.e., remember hits). CR: Correct rejection. NR: Non-recollected target (i.e., know or miss). C) Group-level univariate results: Activity in the left PGp for R and CRs, depicting an effect of both recollection and prior knowledge. Activity was modelled using a standard gamma function. D) Activity in the left PGp for recollection success (R, NR) × prior knowledge (famous, non-famous), while collapsing over encoding duration. R trials showed greater activity than NR trials, and the magnitude of the recollection response increased with prior knowledge. E) Activity in the left PGp for recollection strength via encoding duration (4s, 1s) × prior knowledge (famous, non-famous). R trials with longer encoding durations (i.e., 4s) show greater activity than those that were encoded for 1s.

Next, we sought to characterize the response profile of the left AG to previously studied targets where recollection was successful against when recollection failed (i.e., know response or miss). To this effect, participant-specific parameter estimates from the left PGp were submitted to a 2 (recollection success: recollected, non-recollected) × 2 (prior knowledge: famous, non-famous) repeated measures ANOVA (Figure 3D). Estimates were again collapsed over trials with 1 and 4s encoding duration to increase the number of trials in the famous non-recollected condition. Main effects of recollection success [*F*(1,23) = 43.75, *p* < 0.0001, *partial η^2^* = .66] and prior knowledge [*F*(1,23) = 20.85, *p* = 0.0001, *partial η^2^* = .48] were again observed, while the interaction between both factors failed to reach significance (p > .5). The left AG shows a robust recollection success effect during retrieval, consistent with previous reports (Rugg & King, 2017; Vilberg & Rugg, 2008). Furthermore, the left AG shows greater activity in response to previously studied faces with prior knowledge irrespective of whether recollection succeeds. This again indicates that the effect of prior knowledge is independent of recollection success, modulating the strength of recollection and non-recollection responses in the left AG.

Furthermore, activity in the left AG has been suggested to track recollection strength, or the amount of details recollected, over and above an overall response to recollection (Hutchinson et al., 2014; Rugg & King, 2017). We sought to replicate this effect by testing whether the magnitude of the recollection response in the left AG would increase with longer exposure at study, in line with experimental evidence supporting a recollection strength account (Leiker & Johnson, 2014; Vilberg & Rugg, 2009a; Vilberg & Rugg, 2009b). To this effect, participant-specific parameter estimates from the left PGp were submitted to a 2 (prior knowledge: famous, non-famous) × 2 (encoding duration: 1s, 4s) repeated measures ANOVA, using recollected trials only (Figure 3E). The main effect of prior knowledge [*F*(1,23) = 14.38, *p* = 0.0009, *partial η^2^* = .38] reached significance, but critically, a modest effect of duration was also observed [*F*(1,23) = 4.55, *p* = 0.044, *partial η^2^* = .17] in which faces studied for 4s showed a greater recollection response than those studied for 1s. The interaction between prior knowledge and encoding duration failed to reach significance (p > .7). Overall, we report evidence consistent with notions of recollection strength in the left AG, specifically in the same region that shows sensitivity to prior knowledge more generally.

#### Effect of recollection and prior knowledge across the inferior parietal lobe

We report evidence that the left AG, specifically within the cytoarchitectonically separable region of the PGp, responds to both recollection and prior knowledge within the same subjects. To determine whether the PGp has a unique response profile relative to other regions of the inferior parietal lobe, we extracted participant-specific parameter estimates from each of the 7 cytoarchitectural subregions of the inferior parietal lobe (i.e., PFop, PFt, PFcm, PF, PFm, PGa and PGp; see Caspers et al., 2006, 2013), across both hemispheres, for three contrasts of interest: 1) recollected targets > correctly rejected lures, without prior knowledge (i.e., non-famous), 2) recollected > non-recollected targets, without prior knowledge, and 3) correctly rejected lures with prior knowledge (i.e., famous) > without prior knowledge (i.e., non-famous). Contrasts 1) and 2) isolate recollection effects in the absence of prior knowledge, whereas contrast 3) captures the effect of prior knowledge in the absence of any task-specific recollection. Group-averaged estimates are presented in Figure 4B-D, and significance was determined by two-tailed one-sample t-tests comparing parameter estimates of the sample against 0 for each ROI separately. *P*-values are uncorrected and serve only to provide context for the hypothesis-driven analyses of the left PGp.

**Figure 4.**
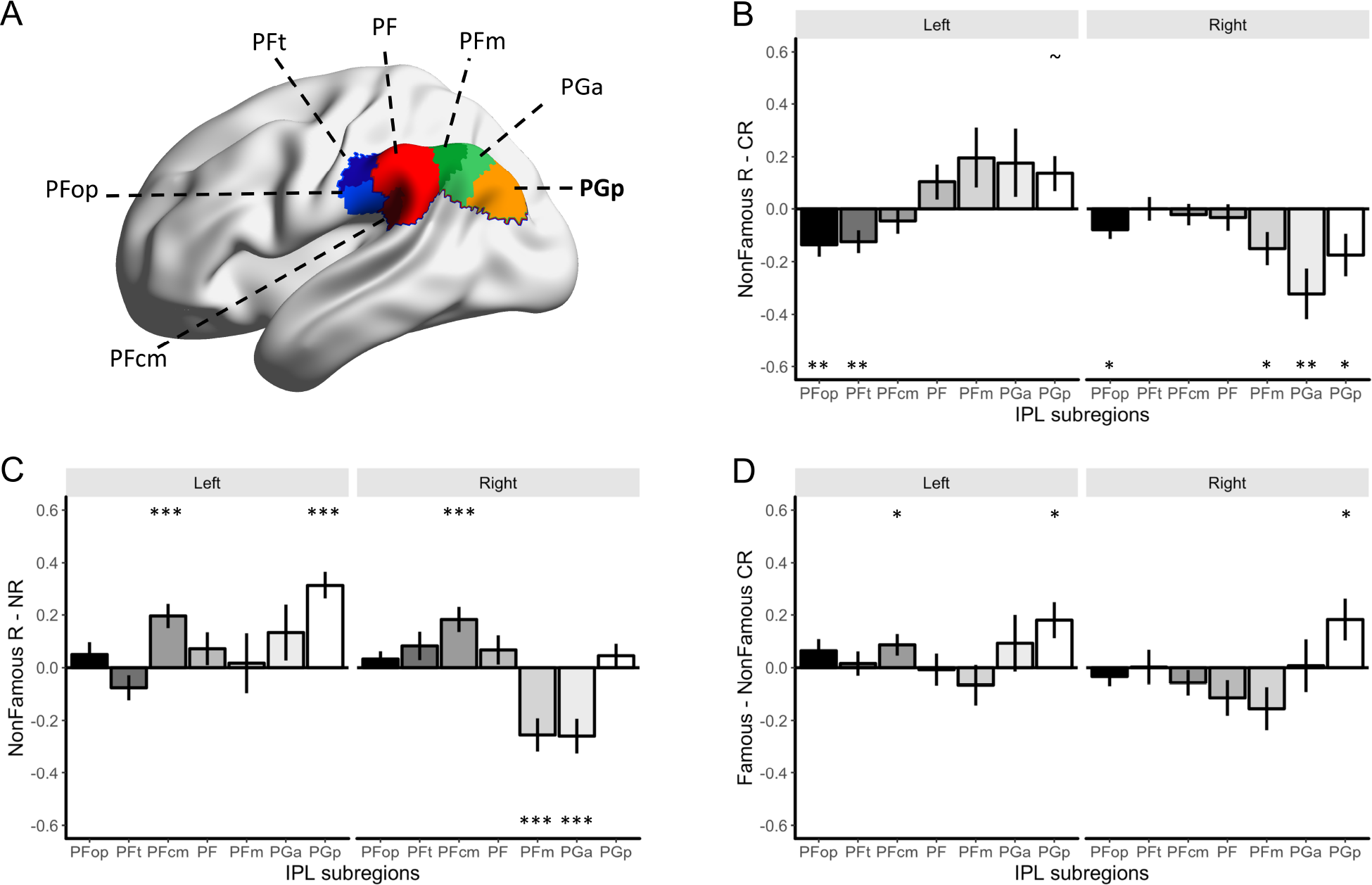
A) Cytoarchitecture-based parcellation of the inferior parietal lobe from Caspers et al., (2008) as implemented in the Julich Histological Atlas. Probability maps were thresholded at 40% and exclusively masked to be non-overlapping. Left hemisphere depicted. B) Group-level contrast examining the effect of R relative to CR in the absence of prior knowledge (i.e., non-famous trials only). Results from contrast are plotted across all subregions in the bilateral inferior parietal lobe. C) Contrast examining the effect of R relative to NR, in the absence of prior knowledge (i.e., non-famous trials only). D) Contrast examining the effect of prior knowledge in the absence of any demands on recollection (i.e., CR only). *** = *p* < .001, ** = *p* < .01, * = *p* < .05, ~ = *p* < .06. Error bars represent standard error of the mean.

For contrast 1), which isolates the effect of recollection relative to correct rejections for trials in the absence of prior knowledge, the left PGp is the only region approaching statistical significance [*t*(23) = 1.34, *p* = 0.059] (Figure 4B). The remaining regions of the bilateral inferior parietal lobe either do not differ from 0 (i.e., Left PFcm, PF, PFm, and PGa; Right PFt, PFcm, and PF; all ps > .1) or show greater activity for correct rejections than recollection [Left PFop: *t*(23) = 3.02, *p* = 0.006; Left PFt: *t*(23) = 2.89, *p* = 0.008; Right PFop: *t*(23) = 2.22, *p* = 0.037; Right PFm: *t*(23) = 2.39, *p* = 0.025; Right PGa: *t*(23) = 3.36, *p* = 0.003; and Right PGp: *t*(23) = 2.17, *p* = 0.04].

For contrast 2), which isolates the effect of recollection relative to non-recollected trials (i.e., know or miss responses) in the absence of prior knowledge, left PGp reaches statistical significance [*t*(23) = 6.09, *p* < 0.0001] (Figure 4C). The left [*t*(23) = 4.27, *p* = 0.0003] and right PFcm [*t*(23) = 3.82, *p* = 0.0009] also show greater activity for recollection success. The remaining regions of the bilateral inferior parietal lobe either do not differ from 0 (i.e., Left PFop, PFt, PF, PFm, and PGa; Right PFop, PFt, PF and PGp; all ps > .1) or show greater activity for recollection failures [Right PFm: *t*(23) = 4.03, *p* = 0.0005; Right PGa: *t*(23) = 3.93, *p* = 0.0007].

For contrast 3), which isolates the effect of prior knowledge in the absence of task-specific recollection, left PGp again reaches statistical significance [*t*(23) = 2.64, *p* = 0.015]. The left PFcm [*t*(23) = 2.11, *p* = 0.045] and the right PGp [*t*(23) = 2.30, *p* = 0.031] also show greater activity for correct rejections with prior knowledge than those without (Figure 4D). The remaining regions of the bilateral inferior parietal lobe largely do not differ from 0 (i.e., Left PFop, PFt, PF, PFm, and PGa; Right PFop, PFt, PFcm, PF, and PGa; all ps > .1), with only the right PFm showing a trend in the opposite direction [*t*(23) = 1.91, *p* = 0.07].

Overall, the left PGp is the only region of the inferior parietal lobule to show increased activity in the hypothesized direction across all three contrasts, highlighting its unique role in responding to both recollection and prior knowledge during retrieval. Interestingly, the right PFm appeared to show a consistently reversed effect for all three contrasts, relative to the left PGp, potentially highlighting this region as sensitive to novelty.

#### Integrating evidence across recollection and incidental prior knowledge

The left PGp, a cytoarchitectonically separable region of the posterior AG, appears to show a unique response to both recollection and prior knowledge relative to other regions of the bilateral inferior parietal lobe. The magnitude of activity in this region also may scale with the amount of information available at the time of recognition, collapsing over idiosyncrasies of a specific episode, as indexed by recollection and more general, relatively semantic information accrued over many similar past experiences, namely, prior knowledge. To directly test this idea of an increase in activity accompanying how much related information across recollection and prior knowledge is available for each trial, we tested for a linear trend at the voxel level examining whether trials with more cumulative lifetime exposure were associated with greater activity. Here, cumulative lifetime exposure is defined as a measure of how much experience a participant had with a given stimulus, in which the degree to which a stimulus is exposed increases with any opportunity a participant had to encounter this stimulus during their lifetime. Critically, this definition of lifetime exposure explicitly incorporates both experience from the experiment itself with pre-experimental knowledge, providing an ideal measure to examine their integration in the brain (see also, Duke, Martin, Bowles, McRae, & Köhler, 2017). For example, a recognized famous face necessarily has more lifetime exposure than a non-famous face by nature of being known pre-experimentally. A face presented for 4s at encoding also has more lifetime exposure than one that was presented for only 1s, as the participant has precisely four times more exposure while studying the former. Similarly, both types of studied faces have more lifetime exposure than a novel lure by virtue of being studied at all. Concretely, a linear increase in brain activity accompanying increased cumulative lifetime exposure was operationalized at the voxel level using the following linear weighted contrast: Famous 4s R > Famous 1s R > Famous correct rejection > Non-famous 4s R > Non-famous 1s R > Non-famous correct rejection (corresponding contrast weights: 5 3 1 −1 −3 −5). This contrast isolates voxels where retrieval activity scales up with the lifetime exposure associated with a given face, summing across experience from both the experimental episode and pre-experimental knowledge. For simplicity, non-recollected trials were excluded from this analysis as their appropriate position on a vector of lifetime exposure is not clear. Whole brain results are presented in Figure 5 for voxels surviving a false discovery rate (FDR) correction of *p* < 0.01. Of the surviving voxels, 342 fall within the left inferior parietal lobe (i.e., combined mask of all 7 subregions), with the vast majority in PGp (43.6%). Percentage of voxels showing a significant linear response in activity to cumulative lifetime exposure from other left IPL subregions are as follows: PF (23.8%), PFcm (13.7%), PGa (9.3%), PFm (1.6%), PFt (1%) and PFop (0.5%). Critically, in addition to the left AG, our voxelwise analysis revealed a distributed set of regions showing a similar linear increase in activity across both recollection and prior knowledge (warm coloured regions in Figure 5; for coordinates and supporting results, see Extended Data: Figure 5-1:4).

**Figure 5.**
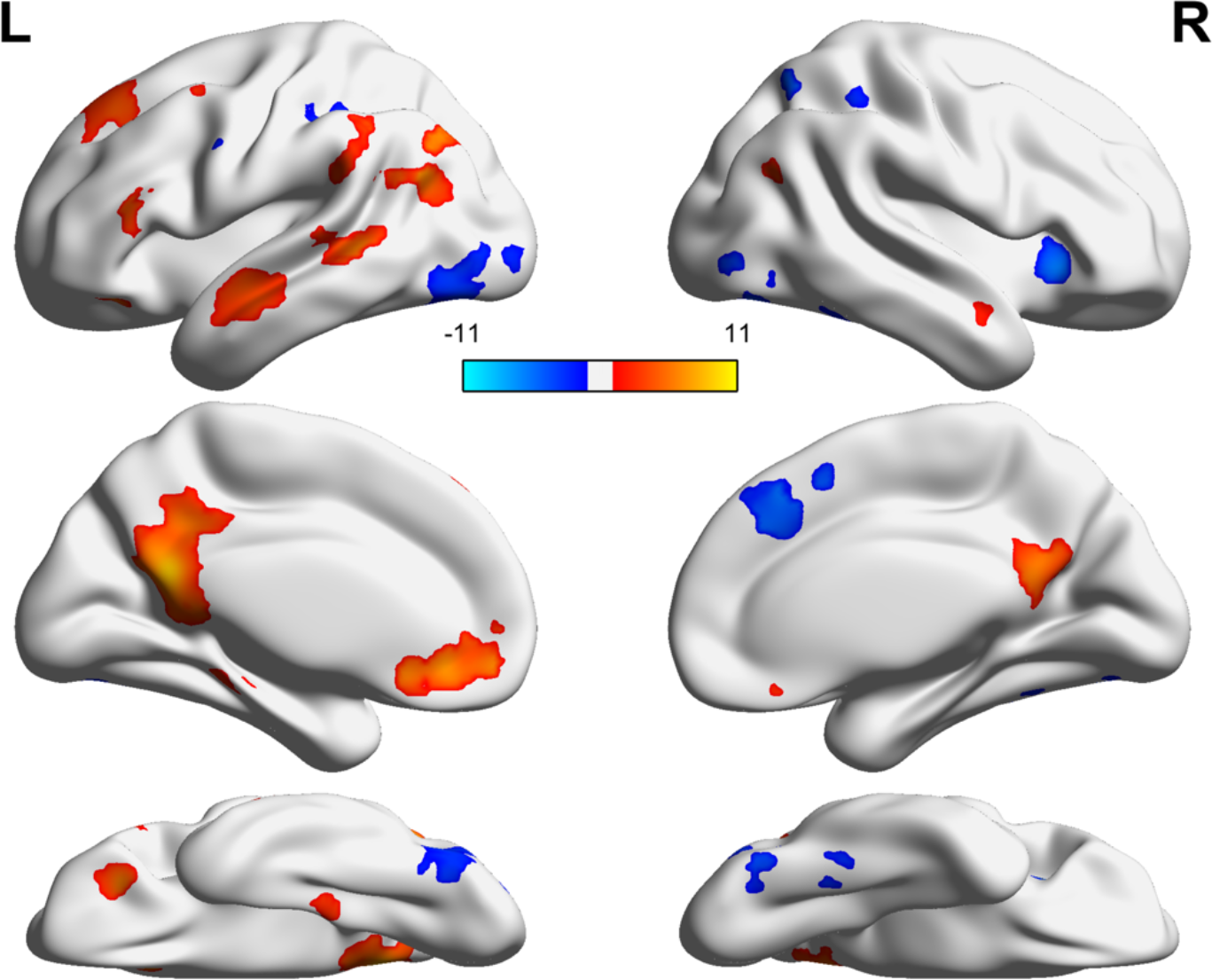
Whole-brain voxelwise linear trend analysis corresponding to cumulative lifetime exposure (i.e., Famous 4s R > Famous 1s R > Famous Correct Rejection > Non-famous 4s R > Non-famous 1s R > Non-famous Correct Rejection; with corresponding contrast weights of: 5 3 1 −1 −3 −5). Warm coloured clusters show a positive linear trend such that magnitude of activity increases with cumulative lifetime exposure. Cool coloured clusters show a negative relationship with cumulative lifetime exposure. Results are thresholded at a false discovery rate (FDR) correction of p < 0.01, with a minimum cluster extent of 20 voxels.

Ventromedial prefrontal cortex, posterior cingulate, precuneus, lateral temporal cortex, right AG, posterior parahippocampal gyrus and the hippocampus, regions commonly known to form the default mode network (DMN), also showed activity scaling positively with increased lifetime exposure. Notably, when reducing the threshold to an FDR correction of p < 0.05, activation extended to the anterior medial temporal lobe, including regions of the perirhinal cortex, as previously observed in other studies of cumulative lifetime exposure (Duke et al., 2017). A separate set of regions, including fusiform gyrus, superior parietal lobe, anterior insula and dorsal anterior cingulate, showed a negative relationship with lifetime exposure suggesting activity in these regions scales up with stimulus novelty.

## DISCUSSION

We provide novel evidence that the left AG, in the vicinity of PGp, is recruited by both experiment-specific recollection and access to pre-experimental semantic knowledge. These independent effects were not generalizable to the broader bilateral inferior parietal lobes. Furthermore, recollection-specific activity in the left AG was heightened for faces with prior knowledge and for those with longer opportunity for study, consistent with accounts of recollection strength modulating retrieval-related activity in this region (Rugg & King, 2017). Critically, the observed strength effect does not appear to be restricted to recollection, as activity in the left AG, alongside a distributed set of regions consistent with the DMN, scaled up with cumulative lifetime exposure across both domains of within-experiment recollection and pre-experimental knowledge. A complete functional account of the left AG during retrieval, therefore, must extend beyond the recollection of specific episodic details to incorporate its concurrent sensitivity to related prior knowledge.

### Angular gyrus, recollection and prior knowledge

Various theoretical accounts have posited why the left AG is involved in our ability to access idiosyncratic past experiences from memory (e.g., Berryhill, 2012; Cabeza, Ciaramelli, & Moscovitch, 2012a; Ciaramelli, Grady, & Moscovitch, 2008; Gilmore, Nelson, & McDermott, 2015; Levy, 2012; Shimamura, 2011; Simons et al., 2010; Vilberg & Rugg, 2008; Wagner et al., 2005), though only recently has its role in representing general semantic knowledge been discussed alongside episodic models (Ramanan et al., 2017; Rugg & King, 2017; Seghier, 2013; Sestieri et al., 2017). Previous meta-analyses have suggested overlap in the AG across semantic and episodic memory (Humphreys & Lambon Ralph, 2014; Kim, 2016), though we provide concrete evidence that activity in the left AG, specifically in the vicinity of PGp, was independently modulated by both recollection and prior knowledge within the same individuals. Furthermore, considering the heterogeneity of the inferior parietal lobe (Caspers et al., 2006, 2013; Nelson et al., 2010), our exploration of the bilateral inferior parietal lobes more broadly revealed the left PGp as the only subregion to show this response profile. These data provide an important link between the semantic and episodic models of AG function: activity in this region cannot be characterized by theories derived from either domain alone. It is also important to note that we observed the left-lateralization commonly reported in neuroimaging studies exploring retrieval effects in the AG (Bellana et al., 2016; Guerin & Miller, 2009), despite using nonverbal stimuli. Although famous faces are associated with names and this verbal information may be activated automatically during retrieval, we also find similar effects for non-famous faces which have no such verbal code. Further work is needed to characterize the nature of this lateralization during memory retrieval and why it occurs.

Few studies have directly probed the AG across episodic and semantic contexts within the same individuals. A recent fMRI study by Bonnici et al. (2016) demonstrated that accessing sensory details from episodic (i.e., studied film clips) and semantic memory (i.e., cued word generation) both recruit aspects of the left AG, suggesting that it may serve as an important junction for the processing of multisensory information. It is unclear whether recruitment of the AG is contingent upon integrating sensory content across distinct modalities. In the present experiment, the left AG was recruited during the correct rejection of novel famous relative to non-famous faces. Novel famous faces in our paradigm should only lead to incidental access of relatively schematic, semantic person knowledge (Bruce & Young, 1986). We cannot preclude the possibility that novel famous faces may also automatically cue associated multimodal details (e.g., voice, scene from a movie) or even episodic memories (Renoult et al., 2012); the task demands, however, do not require any elaboration or explicit integration of these details. Although a multimodal integration account or one based on incidentally cued episodic memories cannot be precluded, there is considerable evidence that the left AG represents abstract, conceptual representations without obvious dependencies on sensory representations or episodic memory. A study by Bonner and colleagues observed activity in the left AG when contrasting abstract words (e.g., doctrine) and pseudowords in the context of a lexical decision task. Similarly, the AG has been reported to show evidence of successful cross-classification across modalities (i.e., using neural representation of the word “apple” to classify a photo of an apple; Fairhall & Caramazza, 2013), consistent with a role in representing abstract, higher-order concepts (Fernandino et al., 2015; Price, Bonner, Peelle, & Grossman, 2015; Price et al., 2016). Therefore, the role of the left AG in integration may not be dependent on processing multisensory information or recalling specific episodes. Instead, we argue that our data provide evidence that recollection-sensitive regions of left AG can also be recruited when accessing related (non-recollected) knowledge, in the absence of explicit demands on multisensory integration or episodic retrieval, consistent with a concurrent role in representing higher-order concepts in semantic memory (Binder et al., 2009; Kim, 2016; Price, 2010).

### Angular gyrus and evidence accumulation

Activity in the left AG can be modulated by the amount of information recollected during retrieval. For example, recollection-responses in the left AG are heightened for trials with longer relative to shorter presentation at encoding (Vilberg & Rugg, 2009a, 2009b), a trend that we replicated (Figure 3E). Similarly, this region has been reported to track stimulus repetition (Gilmore et al., 2015; Guerin & Miller, 2011; Nelson, Arnold, Gilmore, & McDermott, 2013), amount of source information at retrieval (Hutchinson et al., 2014), degree of cortical reinstatement from encoding (Jonker, Dimsdale-Zucker, Ritchey, Clarke & Ranganath, 2018; Kuhl & Chun, 2014; Leiker & Johnson, 2014; Thakral et al., 2017b), and subjective memory strength (Rissman et al., 2016; Thakral et al., 2015). This recent evidence is broadly consistent with the mnemonic accumulator hypothesis (Wagner et al., 2005), which states that activity in the posterior parietal cortex tracks the amount of available evidence for an old response during recognition. Magnitude of activity, or strength of mnemonic evidence, is then compared against a decision criterion ultimately leading to recognition. It is unclear, however, why damage to an evidence accumulator would spare recognition memory performance, as observed in patients with lesions to the inferior parietal lobe (Berryhill, 2012; Rugg & King, 2017). Results from our voxelwise linear trend analysis, however, demonstrate that increased activity in the AG corresponded to a linear vector representing cumulative lifetime exposure, combining recollection strength and prior knowledge on a common scale (also, see Brown, Rissman, Chow, Uncapher & Wagner, 2018). Critically, this was true despite prior knowledge being incidental to recognition decisions in our paradigm. Therefore, the retrieval-related activity in the left AG may track mnemonic evidence beyond what is necessary for recognition, consistent with previous evidence separating AG activity from the decision process itself (Guerin & Miller, 2011). Activity may instead reflect access to a cascade of wide-ranging associations related to the attended target, including contextual details from a past study episodes and broader semantic associations. Therefore, damage to the AG should only affect memory decisions to the degree that this broad ‘associative context’ is necessary. This hypothesis coincides with lesion evidence, where damage to the AG specifically impairs complex memory decisions, such as those probing recollection (Davidson et al., 2008), memory confidence (Hower et al., 2014; Simons et al., 2010), or recall of multimodal details (Ben-Zvi et al., 2015; Ciaramelli et al., 2017).

Overall, we provide evidence that retrieval-related activity in the left AG is sensitive to both experiment-specific recollection and pre-experimental semantic knowledge. In fact, the AG may be particularly suited to integrate recent experiences with prior knowledge. Recent work on temporal receptive windows, or the length of time during which an incoming signal is affected by its previous response history (Hasson, Chen, & Honey, 2015), highlight the long temporal receptive window in the AG (i.e., history-dependence over minutes). This property is ideal for integrating incoming information with extended past experience (Akrami, Kopec, Diamond, & Brody, 2018; Chen, Honey, et al., 2016). Similarly, regions of the DMN (AG, ventromedial prefrontal cortex, posterior cingulate, precuneus, lateral temporal cortex, posterior parahippocampal gyrus and hippocampus) are also characterized by a history-dependence on the order of minutes (Chen, Honey, et al., 2016), highlighting their unique sensitivity to past experience relative to the rest of the neocortex. DMN regions are also the most anatomically segregated from early sensory cortices (Margulies et al., 2016; Murphy et al., 2018) and thus are ideally situated to support complex internal representations removed from immediate sensory input (Mesulam, 1998). Therefore, regions of the DMN, including the left AG, may be modulated by either recollection or prior knowledge when separated experimentally, but are perhaps best characterized by their ability to integrate recent past experience with pre-experimental knowledge in support of complex internal models of an attended target, consistent with their contributions to accessing conceptual knowledge (Binder et al., 1999), episodic memories (Bellana et al., 2017; Rugg & Vilberg, 2013), schemas (Gilboa & Marlatte, 2017; Liu et al., 2016) and cumulative lifetime exposure (Duke et al., 2017). Future research exploring the dissociable functional properties of regions within the DMN is necessary.

## Acknowledgments

The authors would like to thank Rania Mansour for her diligence with stimulus collection, Ali Golestani and Priya Abraham for their assistance at the Toronto Neuroimaging (ToNI) centre, and Marilyne Ziegler for her assistance with Eprime. The authors would also like to thank Bradley Buchsbaum, Michael D. Rugg, Zhong-Xu Liu, Nick Diamond, Vincent Man, Jessica Robin, Tarek Amer and Iva Brunec for many helpful discussions about these data.

## Notes

This research was supported by a Natural Sciences and Engineering Research Council (NSERC) grant to M.M. (no. A8347), a Canadian Institute for Health Research (CIHR) grant to C.L.G. (no. MOP-143311), and scholarships awarded to B.B. from NSERC and the Ontario Graduate Scholarship program.

