## Extended Data for "Recollection and prior knowledge recruit the left angular gyrus during recognition"

**Figure 2-1.** *Prior knowledge facilitates the intentional and incidental formation of durable memories*

To characterize any lasting effects of prior knowledge on recognition memory, we analyzed performance on a surprise delayed recognition memory test presented after the completion of the 8 scanned study-test blocks. First, delayed recognition for intentionally studied targets was examined. Delayed recognition accuracy was defined as the proportion of accurately recognized targets after subtracting the proportion of false alarms on novel lures at delayed recognition (i.e., hits – false alarms), per participant. Only trials that were accurately recognized at immediate recognition contributed to delayed recognition memory analyses. The lasting effects of prior knowledge and encoding duration on delayed recognition accuracy were measured using a 2 (prior knowledge: famous, non-famous) x 2 (encoding duration: 1s, 4s) repeated measures ANOVA (see Figure 2-1-1). A significant main effect of prior knowledge [ $F(1,23) = 33.47$, $p < 0.0001$, *partial* $\eta^2 = .59$ ] and encoding duration [ $F(1,23) = 5.88$, $p = 0.02$, *partial* $\eta^2 = .20$ ] were observed. Furthermore, a significant interaction between prior knowledge x encoding duration also reached significance [ $F(1,23) = 4.53$, $p = 0.04$, *partial* $\eta^2 = .16$ ]. Simple effects using pairwise t-tests demonstrated that the benefit of longer encoding duration on delayed recognition was unique to non-famous faces ($M_{1s} = .692$, $M_{4s} = .745$, $SD_{diff} = 0.156$; $t(23) = 2.61$, $p = .016$, *Cohen's d* = .34) and absent for famous faces ($M_{1s} = .944$, $M_{4s} = .945$, $SD_{diff} = 0.077$; $t(23) = .05$, $p = .96$).

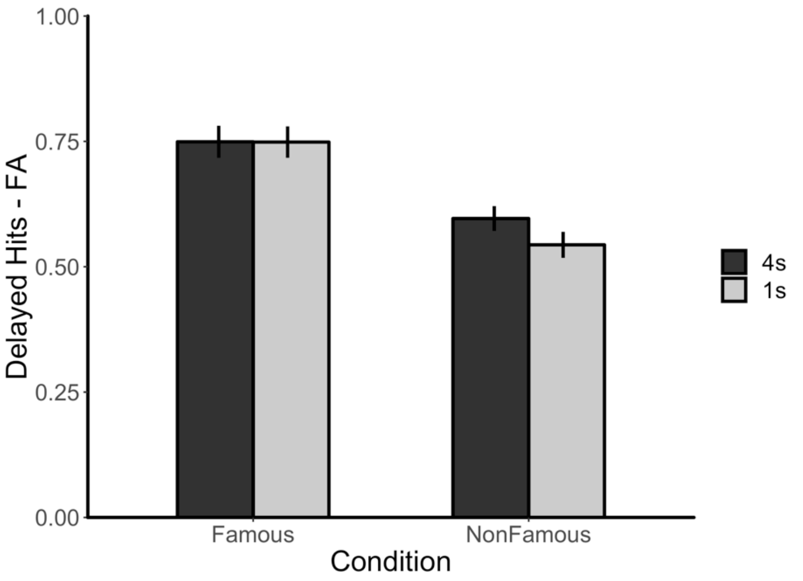

**Figure 2-1-1**. Recognition accuracy (hits – false alarms) from delayed recognition, by prior knowledge (famous, non-famous), and encoding duration (1s, 4s). Error bars represent standard error of the mean.

As all 256 faces from immediate recognition were defined as targets during the delayed recognition memory test, we were able to test delayed recognition for both accurately studied targets and correctly rejected lures from immediate recognition. Correctly rejected lures from immediate recognition were not explicitly studied, and thus delayed recognition of these faces provides a measure of the effect of prior knowledge on incidental encoding. The effect of prior knowledge and encoding type on delayed recognition accuracy was measured using a 2 (prior knowledge: famous, non-famous) x 2 (encoding type: intentional, incidental) within-subects ANOVA (see Figure 2-1-2). Significant main effects of prior knowledge [ $F(1,23) = 66.13$, $p < 0.0001$, *partial* $\eta^2 = .74$ ] and encoding type [ $F(1,23) = 111.5$, $p < 0.0001$, *partial* $\eta^2 = .83$ ] were observed, alongside a significant 2-way interaction between prior knowledge x encoding type [ $F(1,23) = 35.68$, $p = 0.0001$, *partial* $\eta^2 = .61$ ]. Simple effects were conducted using pairwise t-tests comparing the effect of prior knowledge on delayed recognition for intentional and incidental targets separately. Simple effects show that the benefit of prior knowledge on delayed recognition accuracy was present across both encoding contexts but most prominent during incidental ($M_{\text{Fam}}= .618$, $M_{\text{NonFam}}= .260$, $SD_{diff} = 0.200$; $t(23) = 8.76$, $p < .0001$, $d = 1.79$) relative to intentional encoding ($M_{\text{Fam}}= .749$, $M_{\text{NonFam}}= .571$, $SD_{diff} = 0.151$; $t(23) = 5.76$, $p < .0001$, $d = 1.17$). Consistent with our previous report (Bellana, Mansour, Ladyka-Wojcik, Grady & Moscovitch, 2019), this suggests that prior knowledge can facilitate the rapid formation of durable memories automatically, even in the absence of intentional encoding.

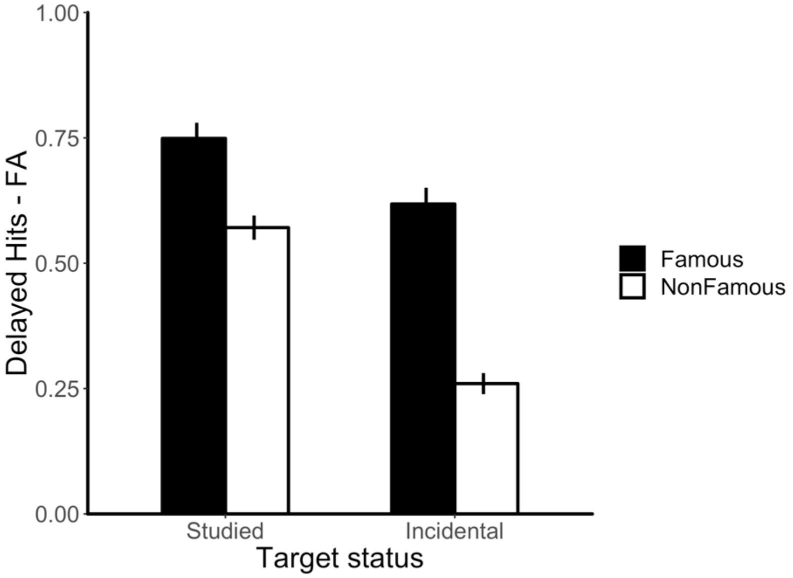

**Figure 2-1-2**. Recognition accuracy (hits – false alarms) from delayed recognition, by prior knowledge (famous, non-famous), and encoding type (intentional, incidental). Error bars represent standard error of the mean.

**Figure 4-1**. *Voxels counts in subregions of the inferior parietal lobe across episodic and semantic memory (Neurosynth)*

| Region of interest | Total number of voxels in ROI (% of IPL) | Episodic voxels (% of IPL) | Semantic voxels (% of IPL) |
|---|---|---|---|
| *Left inferior parietal lobe* | **3254** (100%) | **387** (100%) | **209** (100%) |
| PFop | 300 (9.22%) | 0 (0%) | 0 (0%) |
| PFt | 304 (9.34%) | 0 (0%) | 0 (0%) |
| PFcm | 271 (8.33%) | 0 (0%) | 0 (0%) |
| PF | 818 (25.14%) | 7 (1.81%) | 6 (2.87%) |
| PFm | 339 (10.42%) | 30 (7.75%) | 24 (11.48%) |
| PGa | 420 (12.91%) | 112 (28.94%) | 12 (5.74%) |
| PGp | 802 (24.65%) | 238 (61.5%) | 167 (79.9%) |
| *Right inferior parietal lobe* | **3744** (100%) | **89** (100%) | **72** (100%) |
| PFop | 80 (2.14%) | 0 (0%) | 0 (0%) |
| PFt | 340 (9.08%) | 0 (0%) | 0 (0%) |
| PFcm | 244 (6.52%) | 0 (0%) | 0 (0%) |
| PF | 766 (20.46%) | 0 (0%) | 0 (0%) |
| PFm | 645 (17.23%) | 1 (1.12%) | 0 (0%) |
| PGa | 794 (21.21%) | 17 (19.1%) | 5 (6.94%) |
| PGp | 875 (23.37%) | 71 (79.78%) | 67 (93.06%) |

Total number of 2mm isotropic voxels in final masks for our regions of interest (ROI) are reported above. Percentages are relative to the total number of voxels in the IPL ROI. Masks were calculated from tissue probability maps for the 7 subregions of the inferior parietal lobule (IPL) across both left and right hemispheres from the Jülich Histological Atlas (Eickhoff et al., 2005), thresholded at 40%. These maps were further exclusively masked, from posterior to anterior, to ensure all final ROIs were non-overlapping.

The overlap between our ROIs and Neurosynth (Yarkoni et al., 2011) reverse inference meta-analytic map based on the term “episodic memory” is also reported above. Percentages are relative to the total number of significant voxels in the IPL. This initial exploratory analysis was used to provide an empirical basis for our targeted ROI analysis by highlighting the subregion of the bilateral IPL most commonly activated in the context of episodic memory (http://neurosynth.org/analyses/terms/episodic%20memory/; derived from 270 studies, FDR corrected at $p < 0.01$, 2 mm isotropic voxel resolution).

Considering our concurrent interest in relatively semantic knowledge, we additionally report the overlap between our ROIs and a Neurosynth meta-analysis of the term “semantic memory” (http://neurosynth.org/analyses/terms/semantic%20memory/; derived from 103 studies, FDR corrected at $p < 0.01$, 2 mm isotropic voxel resolution). Again, percentages are relative to the total number of significant voxels in the IPL. Both episodic and semantic meta-analytic maps show the most overlap with the left PGp, consistent with the role of the left AG in accessing specific past episodes via recollection and more general prior knowledge.

**Figure 5-1**. *Voxels counts in subregions of the inferior parietal lobe: Linear trend analysis of cumulative lifetime familiarity*

| Region of interest | Total number of significant voxels (%) |
|---|---|
| *Left Inferior Parietal Lobe* | **342** (100%) |
| PFop | 2 (0.58%) |
| PFt | 4 (1.17%) |
| PFcm | 53 (15.5%) |
| PF | 92 (26.9%) |
| PFm | 6 (1.75%) |
| PGa | 36 (10.53%) |
| PGp | 149 (43.57%) |
| *Right Inferior Parietal Lobe* | **54** (100%) |
| PFop | 0 (0%) |
| PFt | 0 (0%) |
| PFcm | 0 (0%) |
| PF | 0 (0%) |
| PFm | 0 (0%) |
| PGa | 0 (0%) |
| PGp | 54 (100%) |

Total number of 2mm isotropic voxels overlapping between our ROIs the results from the linear trend analysis of cumulative lifetime familiarity are reported above. The statistical map associated with the linear trend analysis was thresholded at a false discovery rate (FDR) correction of $p < 0.01$, with a minimum cluster extent of 20 voxels. Overall, the linear trend of lifetime familiarity is most prominent in the left PGp.

**Figure 5-2.** *Regions modulated by linear exposure from whole-brain analysis (p < 0.01 FDR-corrected, minimum cluster extent = 20 voxels).*

| Region | Hemisphere | x | y | z | beta | # of voxels |
|---|---|---|---|---|---|---|
| Posterior cingulate cortex / Precuneus | Bilateral | -2 | -52 | 20 | 10.59 | 1864 |
| Medial prefrontal cortex | Bilateral | -2 | 54 | -8 | 8.81 | 572 |
| Superior frontal gyrus | L | -24 | 36 | 48 | 5.35 | 515 |
| Lateral temporal cortex / Temporal pole | L | -56 | -8 | -10 | 4.24 | 447 |
| Posterior middle temporal gyrus | L | -60 | -42 | 2 | 6.22 | 391 |
| Posterior angular gyrus | L | -48 | -64 | 28 | 4.08 | 365 |
| Fusiform gyrus (occipital) | L | -30 | -82 | -22 | -4.97 | 319 |
| Dorsal anterior cingulate | Bilateral | 4 | 28 | 46 | -6.81 | 205 |
| Anterior insula | R | 36 | 22 | -4 | -5.97 | 176 |
| Posterior supramarginal gyrus | L | -54 | -46 | 42 | 4.32 | 164 |
| Cerebellum | L | -6 | -78 | -30 | -3.73 | 157 |
| Orbitofrontal cortex | L | -32 | 34 | -14 | 6.06 | 119 |
| Inferior frontal gyrus | L | -46 | 28 | 14 | 4.61 | 89 |
| Fusiform gyrus (occipital) | R | 38 | -74 | -22 | -5.68 | 85 |
| Anterior supramarginal gyrus | L | -46 | -32 | 44 | -2.77 | 74 |
| Anterior middle temporal gyrus | R | 60 | -2 | -16 | 3.24 | 73 |
| Posterior parahippocampal gyrus / posterior hippocampus | L | -20 | -38 | -14 | 3.26 | 71 |
| Temporal occipital fusiform cortex | R | 40 | -46 | -24 | -4.24 | 65 |
| Posterior angular gyrus | R | 48 | -66 | 22 | 4.18 | 54 |
| Occipital pole | L | -30 | -94 | 4 | -2.85 | 50 |
| Middle frontal gyrus | L | -42 | 6 | 54 | 4.83 | 44 |
| Inferior lateral occipital cortex | R | 50 | -68 | -10 | -4.58 | 35 |
| Precentral gyrus | L | -46 | 0 | 36 | -3.74 | 35 |
| Superior parietal lobule | R | 38 | -44 | 50 | -3.06 | 35 |
| Temporal occipital fusiform cortex | R | 32 | -54 | -20 | -3.91 | 34 |
| Superior lateral occipital cortex | R | 28 | -62 | 56 | -4.82 | 30 |
| Inferior lateral occipital cortex | R | 44 | -80 | -2 | -3.63 | 26 |
| Temporal pole | R | 58 | 10 | 0 | 4.33 | 26 |
| Caudate | R | 14 | -4 | 18 | -2.63 | 20 |

**Figure 5-3.** Parametric modulation analysis modelling degree of prior knowledge

Considering the amount of prior knowledge an individual may have about a specific famous face can vary, we ran an additional model implementing a parametric modulation analysis to detect voxels where activity varied according to the degree of prior knowledge associated with each face based on participant-specific 1 (very little) – 5 (a lot) ratings collected post-scan. This model was conducted using AFNI's 3dDeconvolve -stim_times_AM2 option with an identical gamma response model as described in the manuscript, only including additional parametric modulation regressors for: 1) recollected (i.e., remember response) famous trials with 4s exposure at encoding, 2) recollected famous trials with 1s exposure at encoding and 3) correctly rejected new famous trials. Voxelwise beta maps were then averaged across these three trial types, per participant, to provide subject-specific maps of voxels where activity is modulated by degree of prior knowledge across trial types.

Voxelwise parameter estimates for parametric modulation averaged across famous trial types were submitted to one-sample t-tests against 0 (AFNI: 3dttest++). Thresholding with a false discovery rate (FDR) correction of $p < 0.01$ did not reveal any significant clusters. Reducing the threshold to a liberal $p < 0.05$, uncorrected, revealed distributed sets of regions positively or negatively modulated by the specific degree of prior knowledge associated with a famous face. Activity patterns across the parametric modulation and linear trend analysis of cumulative lifetime familiarity were visually comparable, and thus provide a potential extension of the linear trend analysis. Regions of the DMN, including the left AG, are not only sensitive to the presence or absence of prior knowledge, but instead show some evidence consistent with tracking the amount of prior knowledge associated with a given stimulus. Furthermore, regions of the fusiform gyrus, anterior insula and dorsal anterior cingulate show the opposite pattern with increased activity for relatively unfamiliar famous faces than those that are well-known, consistent with their sensitivity to novelty in the linear trend analysis.

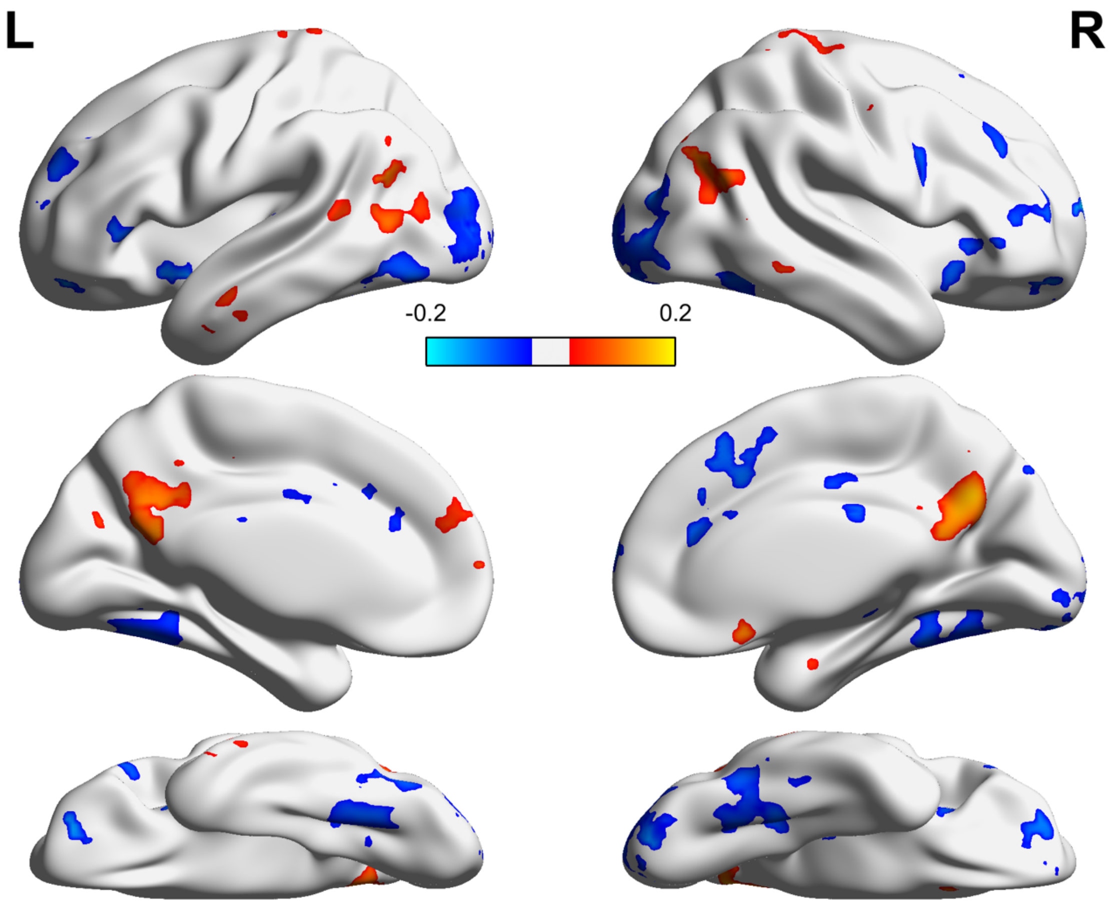

**Figure 5-3-1**. Whole-brain voxelwise parametric modulation analysis on degree of prior knowledge. Warm coloured clusters show a positive relationship between magnitude of activity and fame rating for famous faces. Cool coloured clusters show a negative relationship with fame rating. For visualization purposes, results are thresholded $p < 0.05$, uncorrected, with a minimum cluster extent of 20 voxels.

**Figure 5-4**. *Voxels counts in subregions of the inferior parietal lobe: Parametric modulation analysis on degree of prior knowledge*

| Region of interest | Total number of significant voxels (%) |
|---|---|
| *Left Inferior Parietal Lobe* | **64** (100%) |
| PFop | 0 (0%) |
| PFt | 0 (0%) |
| PFcm | 0 (0%) |
| PF | 0 (0%) |
| PFm | 0 (0%) |
| PGa | 10 (15.63%) |
| PGp | 54 (84.38%) |
| *Right Inferior Parietal Lobe* | **229** (100%) |
| PFop | 0 (0%) |
| PFt | 0 (0%) |
| PFcm | 0 (0%) |
| PF | 0 (0%) |
| PFm | 0 (0%) |
| PGa | 20 (8.73%) |
| PGp | 209 (91.27%) |

Total number of 2mm isotropic voxels overlapping between our ROIs the results from the supplementary parametric modulation analysis are reported above. After scanning, participants rated how much related prior knowledge, on a scale of 1 (very little) – 5 (a lot), they had for each face. These ratings were used in a parametric modulation analysis (for details, see S5 in Supplementary Materials).

The statistical map associated with the linear trend analysis was thresholded at $p < 0.05$, uncorrected, with a minimum cluster extent of 20 voxels (see Figure 5-3-1). Overall, voxels responding to degree of prior knowledge were most prominent in the PGp bilaterally, with the majority in the right hemisphere.
